# PopPhy-CNN: A Phylogenetic Tree Embedded Architecture for Convolution Neural Networks for Metagenomic Data

**DOI:** 10.1101/257931

**Authors:** Derek Reiman, Ahmed A. Metwally, Yang Dai

## Abstract

**Motivation:** Accurate prediction of the host phenotype from a metgenomic sample and identification of the associated bacterial markers are important in metagenomic studies. We introduce PopPhy-CNN, a novel convolutional neural networks (CNN) learning architecture that effectively exploits phylogentic structure in microbial taxa. PopPhy-CNN provides an input format of 2D matrix created by embedding the phylogenetic tree that is populated with the relative abundance of microbial taxa in a metagenomic sample. This conversion empowers CNNs to explore the spatial relationship of the taxonomic annotations on the tree and their quantitative characteristics in metagenomic data.

**Results:** PopPhy-CNN is evaluated using three metagenomic datasets of moderate size. We show the superior performance of PopPhy-CNN compared to random forest, support vector machines, LASSO and a baseline 1D-CNN model constructed with relative abundance microbial feature vectors. In addition, we design a novel scheme of feature extraction from the learned CNN models and demonstrate the improved performance when the extracted features are used to train support vector machines.

**Conclusion:** PopPhy-CNN is a novel deep learning framework for the prediction of host phenotype from metagenomic samples. PopPhy-CNN can efficiently train models and does not require excessive amount of data. PopPhy-CNN facilities not only retrieval of informative microbial taxa from the trained CNN models but also visualization of the taxa on the phynogenetic tree.

**Availability:** Source code is publicly available at https://github.com/derekreiman/PopPhy-CNN

**Supplementary information:** Supplementary data are available at *Bioinformatics* online.

## 1 Introduction

Numerous metagenomic studies of the gut microbiome have linked dysbiosis to many host diseases, e.g., inflammatory bowel disease, diabetes, obesity, cancer, autoimmune diseases, and metabolic disorders [1, 2]. A metagenomic sample is usually described by its microbial taxanomic composition, i.e., the relative abundance of microbial taxa at one of the taxonomic levels (Super-kingdom, Phylum, Class, Order, Family, Genus, and Species), represented as nodes on a phylogenetic tree. The identification of microbial taxa that are associated with the host disease can benefit the early diagnosis, the development of microbial reconstitution (e.g., Probiotic) therapies [3], and the understanding of the disease mechanism [4].

One primary effort on the analysis of the microbiome has been the disease association study and the identification of microbial biomarker signatures for disease prediction. The detection of the associations relies on statistical analyses (parametric or non-parametric) to identify differentially abundant taxa between disease and control groups [5, 6, 7, 8]. However, the association of the individual microbes to a particular type of disease has shown contradictory results [9, 10]. This can be due to various reasons such as the small sample size and the complexity of the diseases.

Alternative approaches using machine learning models, e.g., Random Forest (RF), LASSO and Support Vector Machines (SVMs), and recently, deep neural networks (DNN), demonstrated the potential of developing microbial biomarker signature for the prediction of disease or phenotype of the host [11, 12, 13, 14]. This type of approaches is motivated by the findings that a microbial signature for the host phenotype may be complex, involving simultaneous over- and under-representations of multiple microbial taxa at distinct taxonomic levels and potentially interacting with each other [9, 15]. Varying levels of predictive accuracy have been reported. The performance of the deep learning models is encouraging, owing the ability of deep architectures in identifying potential interactions of microbial taxa for disease prediction [11]. However, the results also raise the skepticism that DNNs may not be suitable learning models due to their requirement of excessive amount of training data, which are impractical in the present metagenomic study. Furthermore, DNNs are often used as black-boxes, making it difficult to extract informative features from the learned models. Therefore, despite the success of DNNs in other biomedical applications [16], it is unclear whether they can outperform the existing models, such as RF, LASSO and SVMs, and whether they can learn a set of informative microbial taxa from metagenomics data.

We have proposed a prototype of a novel architecture for convolution neural networks (CNNs) for the prediction of host phenotype from the microbial taxonomic abundance profiles [17]. CNNs were originally developed based on the visual cortex in images and have been successful in image processing and speech recognition [18]. The major characteristic of a CNN is its ability of generating convolution layers with multiple feature maps that capture the spatial information in training data. However, metagenomic data are represented by relative microbial taxonomic abundance profiles, where taxas can be placed in arbitrary orders. To empower CNNs in metagenomic phenotype prediction, it is important to provide structural input with certain distance metric among the microbial taxas. In our preliminary work, we constructed a phylogenetic tree, a natural structure representing the relationship among the microbial taxa in the profiles [17]. The tree is embedded in a 2D matrix after populating with the observed relative abundance of microbial taxa in each individual profile. In this way, the constructed matrices provide a better spatial and quantitative information in the metagenomic data to CNNs, compared to the vectors of relative microbial taxa abundances in an arbitrary order. Our preliminary analysis has revealed encouraging predictive ability of CNNs based on metagenomic data taken from different parts of body [11, 17].

In the present work, we introduce PopPhy-CNN by extending our prototype CNN learning framework to establish reliable host phenotype prediction models for more complex gut metagenomic data from disease individuals. More specifically, our contribution is summarized as follows.

We investigated the effect of up-sampling in addressing the issue of the moderate datasize in the current metagenomic study. Our experimental results indicates that learning from the original data is sufficient to achieve the maximum performance.

We conducted a comprehensive evaluate of the performance of our CNN model in comparison with other models (RF, LASSO, SVMs) and a baseline 1D CNN using the vector form of relative abundance profiles. We demonstrate the superior performance of our CNN models using three datasets with moderate size: (1) cirrhosis (114 cases vs. 118 controls) [19]; (2) type 2 diabetes (223 cases vs. 217 controls) [20, 21], and (3) obesity (164 cases vs. 89 controls) [22].

We developed a novel procedure to retrieve microbial taxa from the trained CNN models and demonstrated the usefulness of the extracted features for prediction. In addition, we demonstrated a visualization using Cytoscape to facilitate the examination and interpretation of the retrieved taxa on the phylogenetic tree.

## 2 Methods

In this section, we describe the major components of PopPhy-CNN as shown in Fig. 1. First, we show how a microbial taxonomic abundance profile obtained from a sample can be embeded into a matrix format based on the use of a populated phylogenetic tree. Then, we describe our CNN training procedure. Last, the scheme of feature extraction will be presented.

**Fig. 1.**
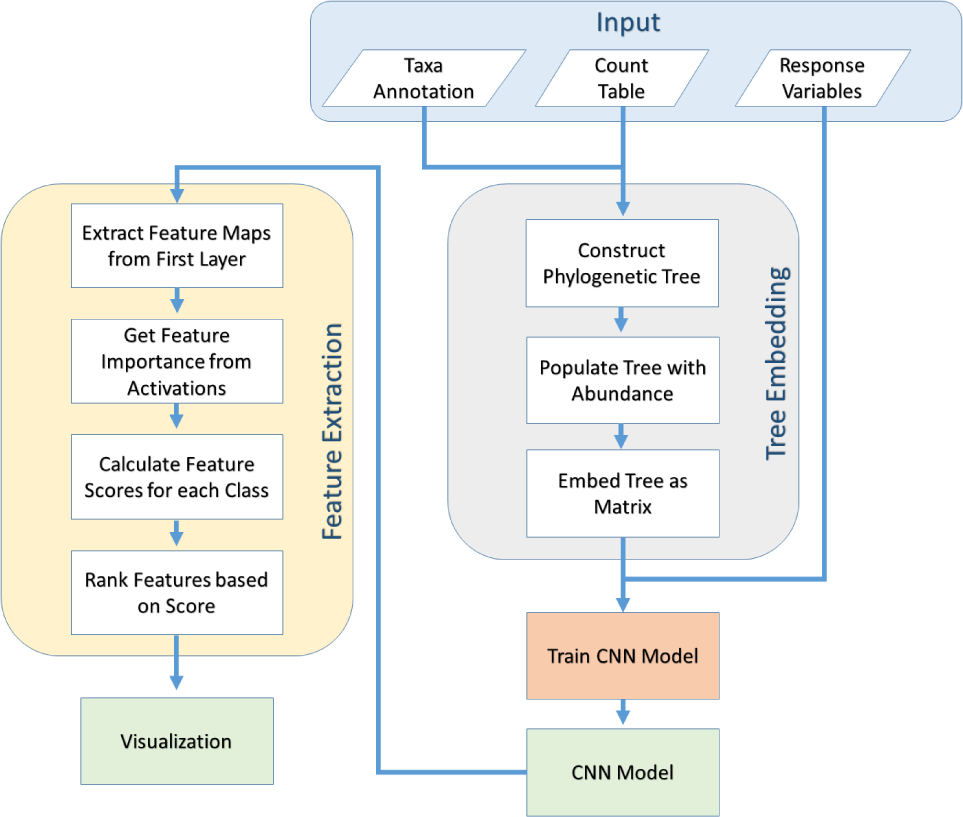
Flowchart of PopPhy-CNN. The taxa annotations and count table are used to create and populate a phylogenetic tree, which is embedded into a matrix format and used to train a CNN model. Features are extracted from the trained model.

### 2.1 Embedding the Phylogenetic Tree

In this section we describe how to transform the microbial taxonomic abundance profiles into a structured data by using a phylogenetic tree. Our method is demonstrated using profile data represented by the Operational Taxonomic Units (OTUs). However, the procedure is applicable to profiles of any level of taxonomic annotation obtained from metagenomic study. Fig. 2 shows an example of converting an OTU vector into an input matrix for CNNs.

**Fig. 2.**
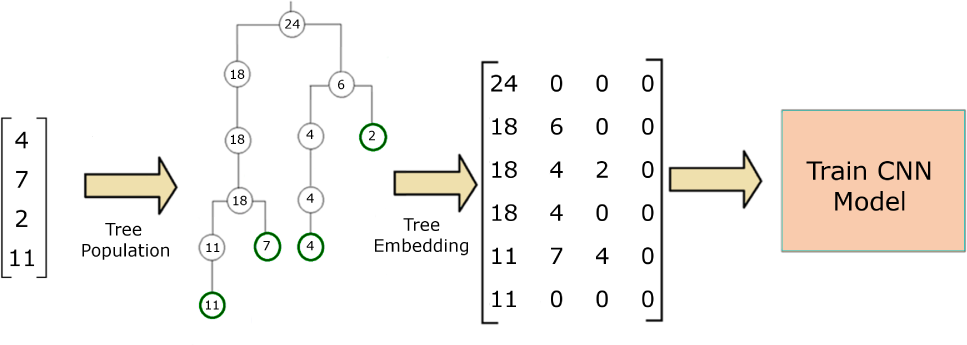
Example of populating and embedding phylogenetic tree. The OTU labels in the vector of an abundance profile are used to construct a phylogenetic tree. The abundance data are used to populate the tree, which is then embedded into a matrix to be used as an input for the CNN.

A phylogenetic tree captures similarity information among OTUs. It can be constructed by comparing the microbial genomes based on multiple sequence alignment and organizing similar taxa into clades. The similarity between taxa is represented by their closeness in the tree. In our work PhyloT [24] and iTol [25] were used to create and visualize the phylogenetic tree. The phylogenetic tree is structured using ancestral nodes from both taxonomic groups and subgroups with no defined distances between nodes. This leads to a tree with more than just seven layers and with branches of variable length. In our work, a constant distance of one between nodes is assumed. This allows for the distance between any two nodes to be determined by the number of nodes between them.

Since CNNs are very successful in image processing where inputs are a multi-dimensional matrix, we need to embed the tree into a matrix format that contains meaningful similarity information both vertically through the rows and horizontally through the columns. We began by combining the OTU abundances and phylogenetic tree by assigning the nodes corresponding to our OTUs their respective abundance value. This is followed by populating the rest of the tree where a parent node’s abundance is the sum of its children’s abundances. This is preformed from the bottom upwards to the root node, which is populated with the sum of the abundance from all organisms found in the community. This procedure is described in **Algorithm 1**.

**Algorithm 1.**
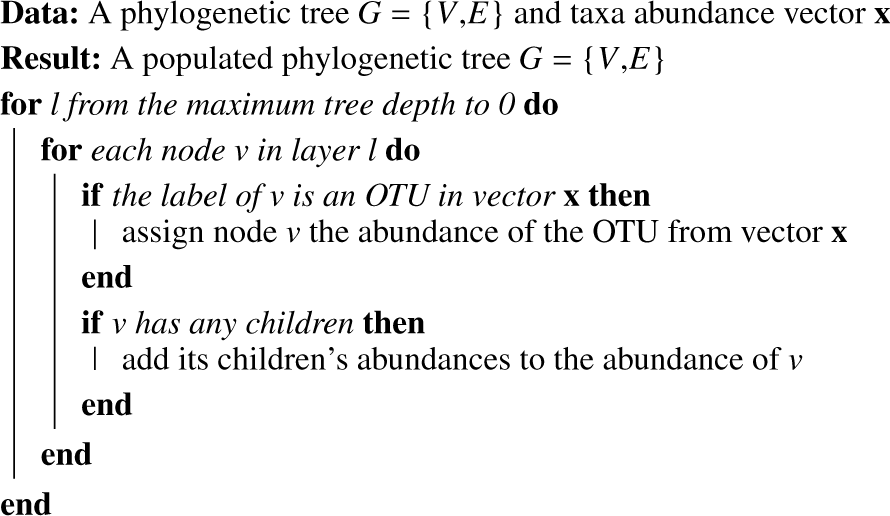
Tree Population

After being populated with abundance data, the tree now contains data representing both abundance as well as phylogenetic similarity. The next step is converting the tree into a matrix format in such a way as to preserve both aspects. To do this, we start by placing the root in the top left corner of the matrix. Then moving down the tree layer by layer, the matrix was filled by taking the set of the child nodes in the order that they appear in the tree from left to right and padding with zeroes to create a twodimensional matrix. The embedding creates a dense section of data that captures both the hierarchy of the tree in its rows as well as similarity between tree branches within its columns. The matrix is then used as the input for the CNN. The algorithm for matrix embedding is described in **Algorithm 2**.

**Algorithm 2.**
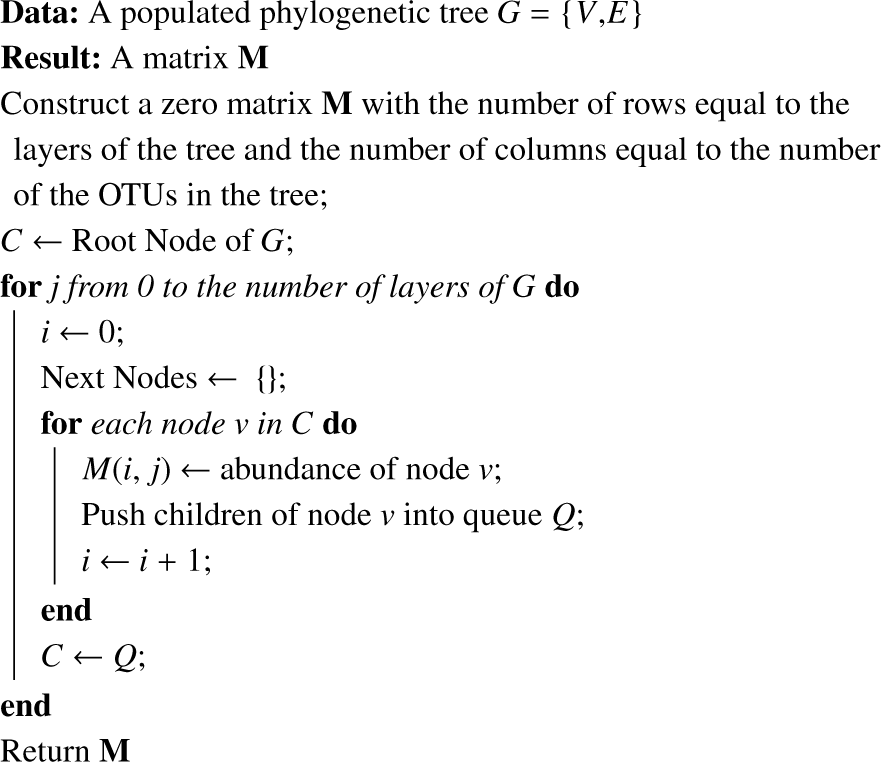
Tree Embedding

### 2.2 Architecture of Convolution Neural Network

Standard CNNs are composed of multiple convolutional layers, which are usually followed by at least one fully connected layer. Each convolutional layer is composed of multiple kernels, each of which transforms the input of that layer into a feature map of velocities through a convolutional operation. For a given kernel *k* with weights *W*^(*k*)^ of size *m* x *n* and input *X*, the velocity of point (*i, j*) is calculated as:

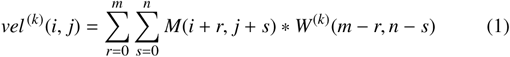

The feature maps composed of these velocities are then passed through a non-linear activation function and subsampled through max or mean pooling to give a matrix of activations.

The CNN architecture used in this study consists of three convolutional layers, two fully connected layers, and a single output layer. Each convolutional layer contained 64 filters and used the rectified linear unit (ReLU) activation function, which sets all negative values to zero while not changing the positive values. Max-pooling was then used for subsampling. The fully connected layers each contained 1,024 neurons and also used the ReLU activation function. The ReLU activation function was chosen since it has been shown to speed up the training time while still maintaining the non-linearity provided by other activation functions [26].

There is a single output neuron for each class in the dataset. Since our datasets all had binary response variables, the output layer contained two neurons. We applied the softmax activation function to the output layer. The softmax activation is a convex function that creates a probability distribution over the number of possible outputs. Given a set of values, {*y*_1_, *y*_2_,…,*y*_J_}, the softmax of any value *y*_*i*_ from the set is calculated as: Softmax(*y*_*i*_) = 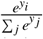 When considering our outputs in the network, the output node with the highest softmax value is selected as the winner and is returned as the predicted output for the given input.

We chose to use the log-likelihood cost function for our model. In order to prevent any bias from class imbalance, we applied a penalty weight that is determined by the total number of samples divided by the number of samples in a given class. This would scale the cost in a way such that samples of less frequent classes were scaled higher, balancing out any bias from the more frequent classes. We then regularized our model by adding an *l*2 penalization to the weights in order to prevent large weight values. Therefore, our final model was trained using the cost function:

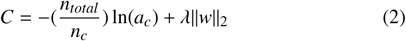

Where given an input whose true label is *c, a*_*c*_ is the output activation for class *c, n*_*total*_ is the total number of samples, *n*_*c*_ is the number of samples for class c, and *λ* is the regularization parameter to penalize the weights.

To help prevent overfitting, we further applied dropout to the fully connected layers [27]. The dropout method works by randomly selecting nodes within the hidden layers and temporarily removing them from the training, preventing both feed-forward information from that node as well as feedback information from back-propagation. This allows the network to train subnetworks to reach the desired output, creating multiple paths to predict the correct output. An overview of the CNN architecture used can be seen in Fig. 3.

**Fig. 3.**
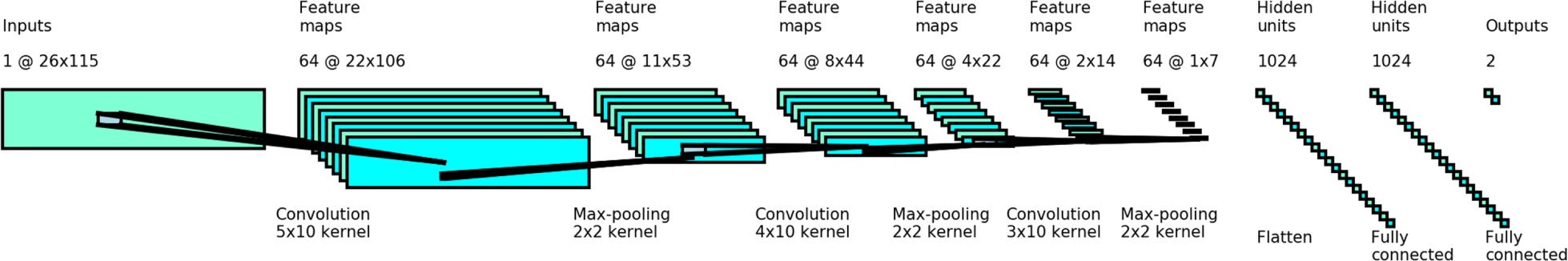
Architecture of PopPhy-CNN. The convolutional neural network is composed of three convolutional layers. Each layer contains 64 kernels and uses max-pooling and ReLU as the activation function. The output from the last convolutional layer is passed to two fully connected layers of 1024 neurons and then finally to a softmax output layer with 2 neurons.

### 2.3 Extraction of the informative features from learned CNN models

One of the biggest criticisms of deep learning models is the lack of interpretability. In our work, we attempt to push past the black-box of the CNN model and try to extract which features the model found to be important for the prediction. A previous study has shown that using feature maps captured by CNN models as features for other machine learning models (i.e., RF and SVMs) yielded better results than using the raw features [28]. This led us to believe that the activations found within the feature maps could be used to evaluate the raw features. Even though deeper layers yielded better features in the previous study, the loss of resolution through subsampling and extra layers of nonlinear transformations could impact the interpretability of the activations. Therefore, we focused on the post analysis of the feature map activations in the first convolutional layer prior to subsampling and the application of the ReLU activation function in order to evaluate which positions in the input contribute most to the highest activations in the learned CNNs.

To do this, we first calculated all of the feature maps for each sample in the training set using the weights from the first convolutional layer. Next we looked at the feature maps generated by a single kernel, *k*, across all the samples for a specific class, *c*. For each of these feature maps, we took note of the positions for a number of maximum values specified by a given hyper-parameter, *θ*_1_ and kept track of the frequency that each position was found in the top portion of velocities. We then selected the top *θ*2 most frequent positions that were found in the maximum velocities. For each velocity selected, we traced its location in the feature map (*u, v*) back to the submatrix of the input *M* from which it was calculated. We call this matrix *R* our reference window. More specifically, given a kernel *k* with weights *W*^(*k*)^ with dimensions *r* x s,

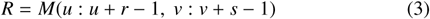

Within the reference window, every position (*i, j*) is equivalent to some node *v* from the phylogenetic tree with an OTU label, *f*. We calculate the importance of each feature *f* given the reference window *R* for sample *S* >as its proportion of the velocity.

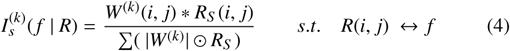

Here *k* is our current kernel with weights *W*^(*k*)^ which have been flipped to account for the convolution function; the summation is over all positions in *R_S_*. The absolute value of the weights in the denominator was used in order to handle any case where the contribution of one large positive component and one large negative component can give rise to an velocity that is much smaller than its components. This could lead to sporadic scaling of importance values which would be hard to interpret. By using the absolute value of the weights, we created an upper boundary of 1 and a lower boundary of -1, and the sum the absolute value of all importance values within a given reference window will sum to 1.

Within a single reference window, some taxa may have be found important in a small subset of the samples but may not be important considering all of the samples. In order to capture only the taxa which were consistently found important, we calculated the mean importance value of a feature *f* across all samples in class *c* given a single reference window *R* and kernel *k*.

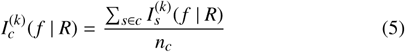

Since the reference windows of different velocities may overlap, it is possible for a single feature to have multiple importance values using the same kernel. A feature may also be found to be important by multiple kernels. This can lead to multiple importance values for a single feature. In order to handle this problem, we selected the importance of *f* to be the maximum over all reference windows containing *f* and over all kernels, *k*

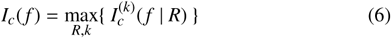

Lastly, we assigned a score for a feature from the perspective of class *c* as the difference of the feature importance using all the samples within the class and the feature importance using all the samples not in the class. Given only two classes, the scores will be the same values with opposite sign. Despite that, we designed our method to be able to handle scenarios where there are more than two classes.

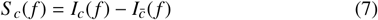

From these scores we created a list of feature scores for each class, allowing the analysis of feature importance from the perspective of different classes that can then be ranked. If an OTU was not found in any of the kernels, it is ranked at the bottom the feature list. The pseudo-code for feature extraction is described in **Algorithm 3**.

## 3 Results

### 3.1 Datasets

In our experiment, we used three publicly available datasets contained within the MetAML package [14]: cirrhosis, type 2 diabetes (T2D), and obesity. They were selected due to the varying difficulty among them. The cirrhosis dataset was taken from a study of 114 subjects with cirrhosis and 118 healthy patients [19]. The type 2 diabetes dataset was a combination of two studies [20, 21] yielding a total of 223 patients with type 2 diabetes and 223 healthy patients. The obesity dataset comes from a study of 292 individuals of which 89 individuals with a BMI lower than 25 *kg/m*^*2*^ were studied against 164 individuals with a BMI greater than 30 *kg/m*^*2*^ [22]. Each of these datasets was generated using whole metagenome shotgun (WMS) sequencing. In the MetAML study [14], the OTUs for each dataset were assigned by MetaPhlAn2, which selects OTUs based on the read coverage of clade-specific markers and then estimates their relative abundance [23].

The OTUs in each dataset were aggregated at the genus level using MetAML’s built-in filter method. The genus level was chosen since PhyloT was not able recognize many OTUs at the species level. For any OTU which was specified as "unclassified" at the genus level, we instead created a feature containing the label for the family level name of that OTU. To handle this OTU when populating the tree, we added this feature to the sum of its children. A summary of the three datasets used in our evaluation is shown in Table 1.

**Algorithm 3.**
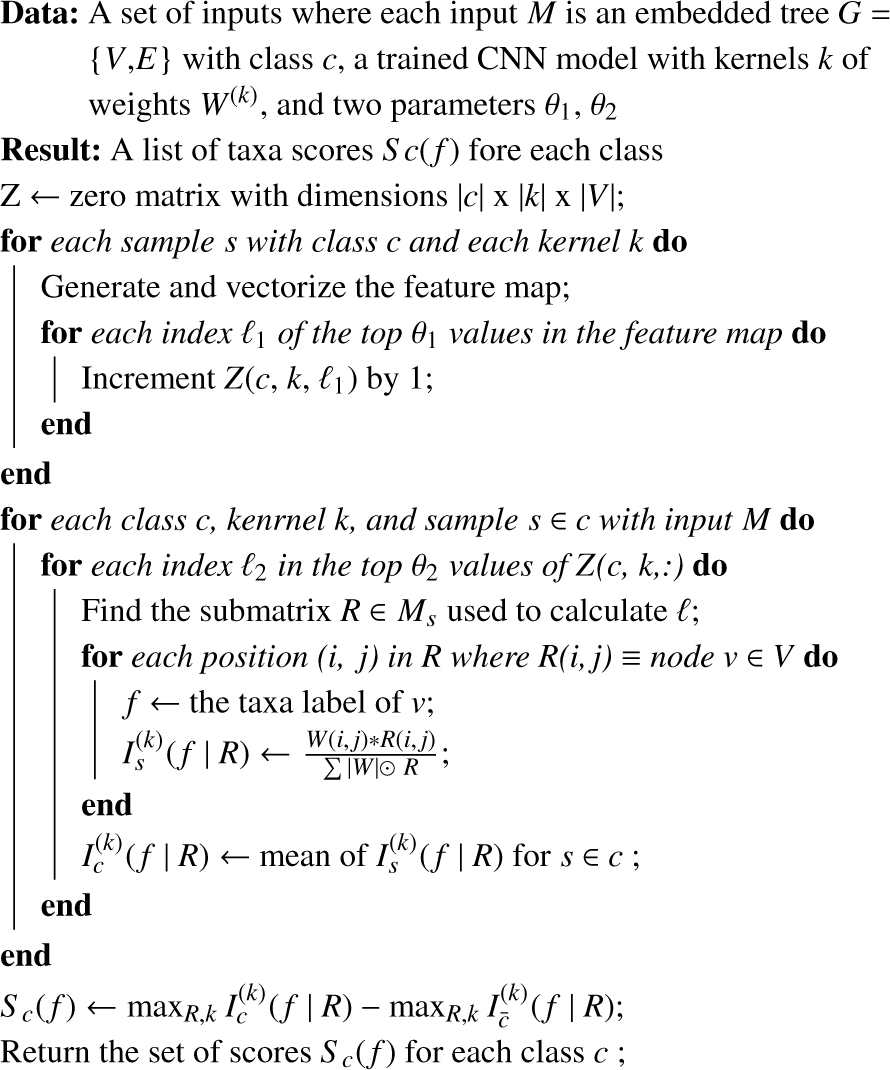
CNN Feature Extraction

**Table 1.** Summary of datasets

|  | # of Case | # of Control | # of Features |
| --- | --- | --- | --- |
| cirrhosis | 114 | 118 | 184 |
| T2D | 223 | 227 | 214 |
| obesity | 164 | 89 | 181 |

### 3.2 Model Evaluation

Since CNNs are often believed to require very large training sets for training, and the three datasets described above are relatively small, we first tried to increase our sample size by re-sampling and adding noise to each new sample. However, we noticed a slight decrease in prediction performance when comparing models trained on the original dataset versus models trained using up-sampling. Therefore, we evaluated PopPhy-CNN without re-sampling.

PopPhy-CNN was benchmarked against RF, SVM, LASSO methods, and a 1D-CNN model taking the abundance vectors as inputs. The 1D-CNN model serves as the baseline to evaluate if the addition of phylogenetic information improves the prediction in CNN. Each model underwent 10 times 10-fold cross validation, using the same partitions across all methods, with an exception for SVM where the data were min-max normalized before training and testing. We found that without this normalization, the SVM models would not converge.

RF, SVM, and LASSO were trained using Python’s scikit package. In RF training a maximum of 500 trees was set and all other parameters were left as the default. The SVMs were trained using a grid search 5-fold cross validation over the linear and Gaussian kernels with an exhaustive search using the set 1, 10, 100, 1000 for error terms and the set 0.001, 0.0001 for γ values in Gaussian kernels. The LASSO model was trained using iterative fitting of the error term α using the set of 50 numbers from 10^−4^ to 10^−0 5^ that were spaced evenly on a log-scale. The best model parameters were again evaluated using 5-fold cross validation.

In order to train the CNN models under cross validation, the network was trained for a number of epochs until the network was deemed to be overfitting the training data. We observed that for each dataset, this usually occurred between 300 and 400 epochs and was determined when the training accuracy was greater than 0.98. However, in order to capture the best model during the training process we calculated AUC, the area under the receiver operating characteristic curve (ROC) after each epoch. If the new AUC value was greater than the previous best, the maximum was updated and the network was saved. This allowed us to retain the best model during training before the model began to overfit the data. Each network was trained using stochastic gradient descent with a dropout rate of 0.5, *λ* of 0.1, and a learning rate of 0.001.

Our evaluation is summarized in Fig. 4. In all three datasets, the 2D-CNN, i.e., PopPhy-CNN models outperform the other methods. The 1D-CNN model is as competitive as RF, which performs better than SVM and LASSO. Our results shows that not only is the PopPhy-CNN able to outperform the other state-of-the-art methods, despite having a relatively small datasets, but also that the embedding into a matrix to spatially capture the phylogenetic information also improved the performance. PopPhy-CNN achieves a mean AUC value of 0.940 in the cirrhosis dataset, 0.753 in the T2D dataset, and 0.676 in the obesity dataset.

**Fig. 4.**
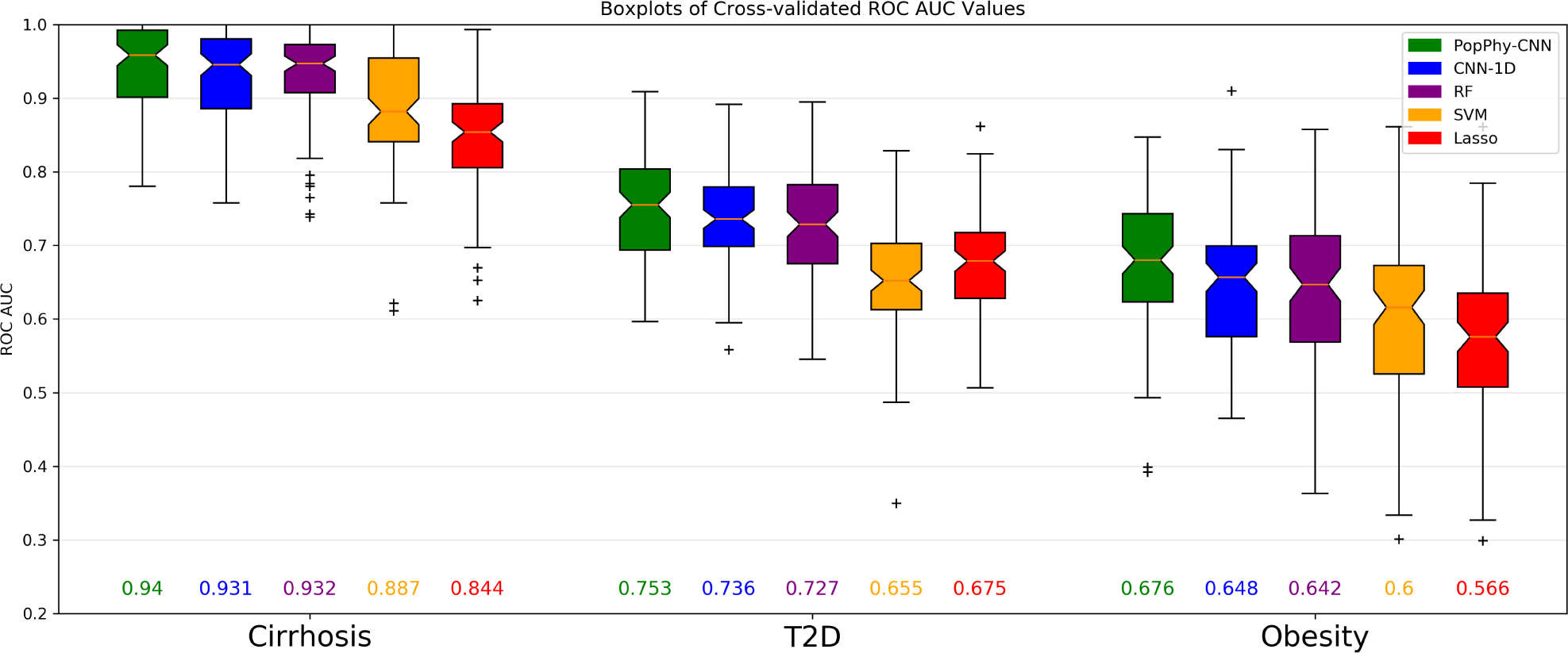
Boxplots comparing SVM, RF, LASSO, 1D-CNN, and PopPhy-CNN across the three datasets: cirrhosis, type 2 diabetes, and obesity. The boxplots were generated using the 100 AUC values for each method after training 10 times 10-fold cross validation. The numbers below each boxplot are the mean cross validated AUC values.

When comparing the 2D-CNN and the 1D-CNN, we observed that there was more improvement in the more difficult datasets. The lowest improvement of 0.009 in AUC values was observed in the cirrhosis dataset, while a larger improvement of 0.017 in the more challenging T2D dataset. In the most difficult dataset, obesity, we saw an improvement of 0.028. Furthermore, we observe the CNN models appear to be more stable than the previous state-of-the-art methods, having a smaller standard deviation of AUC values as well as fewer outliers. In summary, PopPhy-CNN outperforms the other machine models without requiring a large amount of training data.

### 3.3 Extracted Features

Feature rankings for each dataset were generated following the procedure outlined in the Methods section. In the results shown, we did not filter only the top activations, but rather we used all of the activations in the feature maps. We did this because we believe that Θ_1_ and θ_2_ are exploratory hyper-parameters and need to be tuned for each dataset. In order to evaluate the feature extraction fairly, we felt it was best to forgo this hyper-parameter in our comparisons. Since the CNN was trained using 10 times 10-fold cross validation, each feature had 100 sets of rankings. To evaluate the features, we ordered them by the median rank across the 100 models where a lower value represents a higher rank. The top ten ranked features from the cirrhosis dataset without filtering can be found in Fig. 5. Fig. S1 shows the same results after filtering the top 10% of maximum activations and frequencies.

**Fig. 5.**
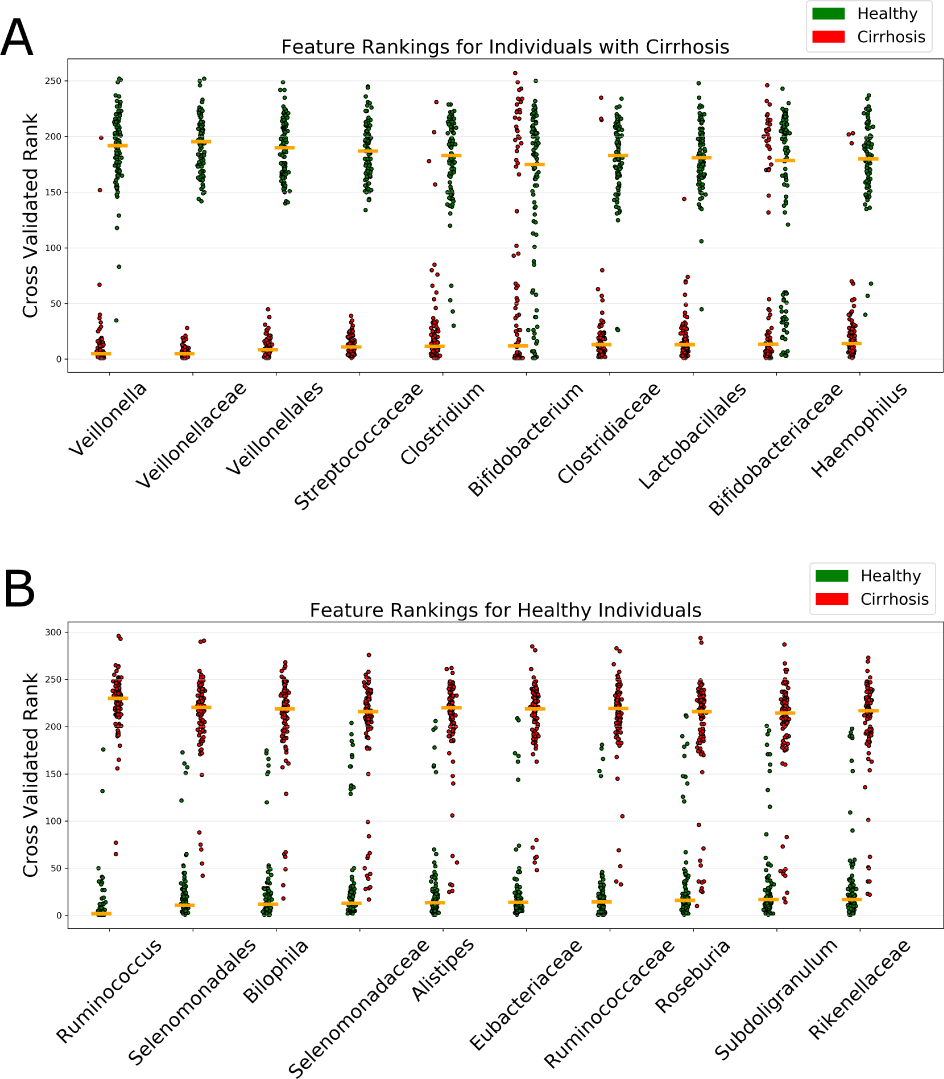
Feature rankings for cirrhosis dataset without filtering for (A) the top ten features found in cirrhosis patients and (B) the top ten features found in healthy patients. Green points represent the rank in the healthy patients and red points represent the ranks in the patients with cirrhosis. Each point represents that feature’s rank based on ***I**c*(*f*) in a given cross-validated model. Features are ranked along the x-axis by the median of their importance across the 100 trained models.

The rankings of the features in the diseased class separated more cleanly then the rankings of the healthy features (Fig. 5 (A)). We found that *Veillonella, Streptococcus, Haemophilus*, and *Lactobacillus* were found important in the cirrhosis subjects. In healthy individuals, *Alistipes, Rumminococcus, Roseburia*, and *Eubacterium* were found to be the most important feature, although the score was not as significant as the those found in the Cirrhosis subjects.

We compared the features found based on our procedure to those reported in the original study [19] and observed that PopPhy-CNN found all significant OTUs found in the cirrhosis patients as well as additional ones that have not been reported. The model also captured most of the features found in the healthy patients. The original study showed that the difference in the OTUs found in the diseased patients were more significant, which supports our finding that the features identified in the disease patients in our study are more discriminatory than the features found in the healthy patients.

The features for the T2D dataset can be found in Fig. S3. We analyzed the features both by filtering the top 10% activations and locations as well as using all the features. Despite the difference in filtering, the top features between the two analyses were similar. The PopPhy-CNN model identified *Lactobacillus, Coprobacillus*, and *Megasphaera* as the strongest discriminitive features found in patients with type 2 diabetes. It also identified *Haemophilus, Streptococcus, Faecalibacterium*, and *Roseburia* as predictive of the healthy patients in this dataset. The rankings of the extracted features in the Obesity dataset (Fig. S4) have much larger differences between when the maximums were filtered and when all velocities of the feature map were used. We did notice that both analyses found *Ruminococcus, Prevotella*, and *Escherichia* in the healthy patients and only *Verrucomicrobia* was common between the two analyses when looking at the features in the lean patients.

### 3.4 Visualization of Extracted Features

We used Cytoscape to visualize the phylogenetic tree and annotate the nodes and edges based on the calculated importance scores, *S*_*c*_(*f*). For this, for each feature we subtracted the score from the healthy samples from the score from the diseased samples, creating a scale in which positive scores constitute important features found in diseased patients, negative scores constitute important features found in healthy patients, with zero representing unimportant features. The feature extraction not only capture features at the OTU level, but the ancestral nodes as well, which can be observed from the visualized the phylogenetic tree. The phylogenetic tree created from the cirrhosis dataset can be seen in Fig. 6. A color gradient was applied to the nodes and edges. For nodes in the tree, healthy features are colored green and disease features are colored red, with yellow constituting unimportant features. The edges are colored in a similar gradient based on the average score of the connected nodes.

**Fig. 6.**
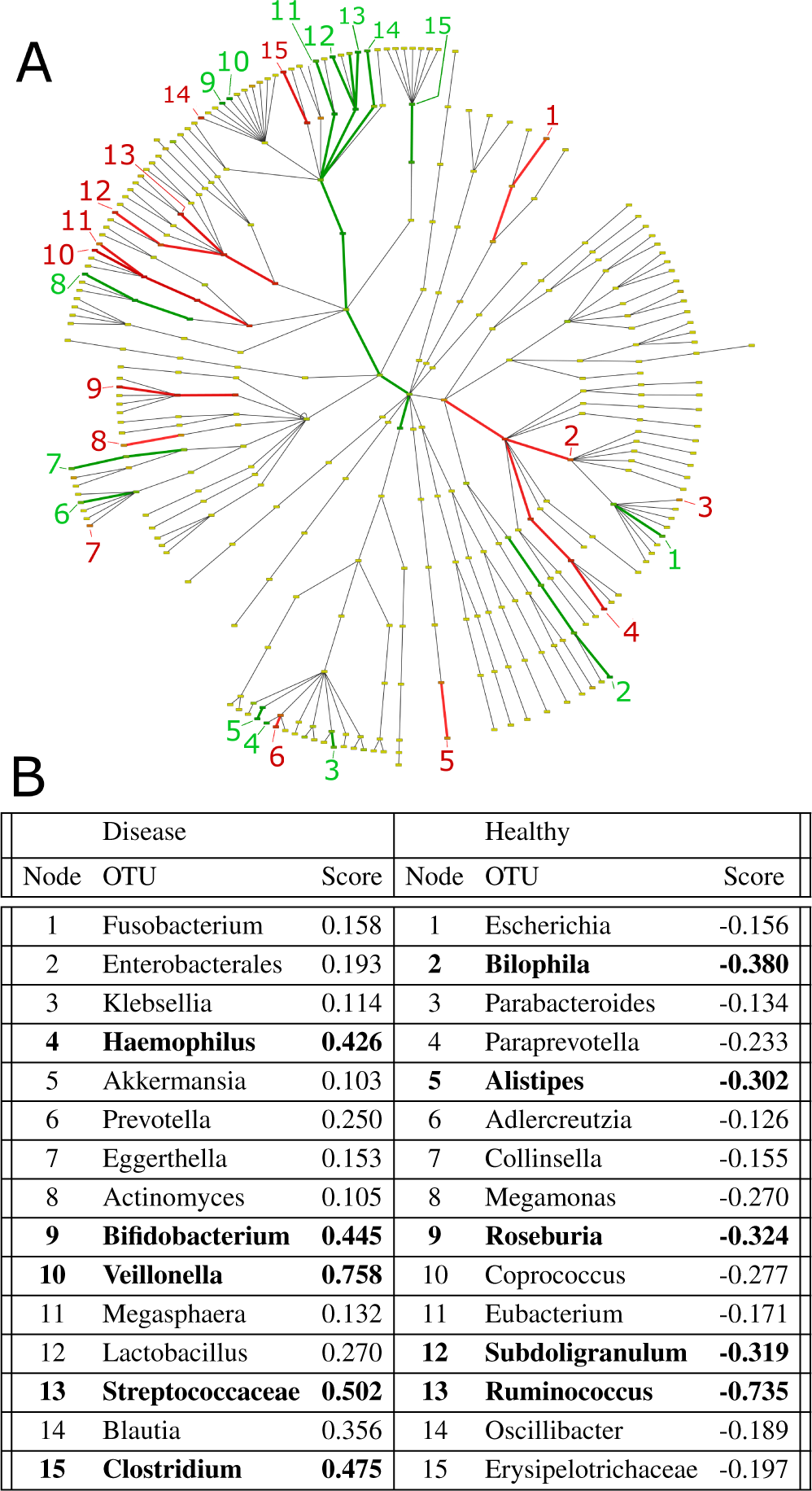
Visualization of the cirrhosis features found by PopPhy-CNN. (A) An annotated phylogenetic tree from the cirrhosis dataset showing sub-trees found important in the disease patients (red) as well as the healthy patients(green). (B) The table highlights features at the bottom of a subtree and whose score’s magnitude is greater than 0.1.

We then used Cytoscape to visualize the phylogenetic tree and annotate the nodes and edges based on the calculated importance scores. For this, for each feature we subtracted the score from the healthy samples from the score from the diseased samples, creating a scale in which positive scores constitute important features found in diseased patients, negative scores constitute important features found in healthy patients, and zero represents unimportant features. The feature extraction was not only able to capture these features, but the ancestral nodes as well, which can be observed from the visualized the phylogenetic tree. The phylogenetic tree created from the Cirrhosis dataset can be seen in Fig. 6. A color gradient was applied to the nodes and edges. For nodes in the tree, healthy features are colored green and disease features are colored red, with yellow constituting unimportant features. The edges are colored in a similar gradient based on the average score of the connected nodes. A gradient was also applied to the edge thickness in order to better visualize the difference in scores. This visualization can facilitate the interpretation of extracted features in the context of the phynogenetic tree.

### 3.5 Evaluation of Extracted Features for Prediction

We compared the extracted top 20 features found in PopPhy-CNN, RF, and LASSO for each dataset. In PopPhy-CNN’s extracted features, about half of the features were not from the original OTUs, indicating the ability of PopPhy-CNN in finding important hierarchical combinations. However, for comparison with the other methods, the overlap of the top 20 OTUs found in the original input vectors was plotted in Fig. 7(A). We observed that the overlap was not large and it becomes smaller in the harder datasets.

**Fig. 7.**
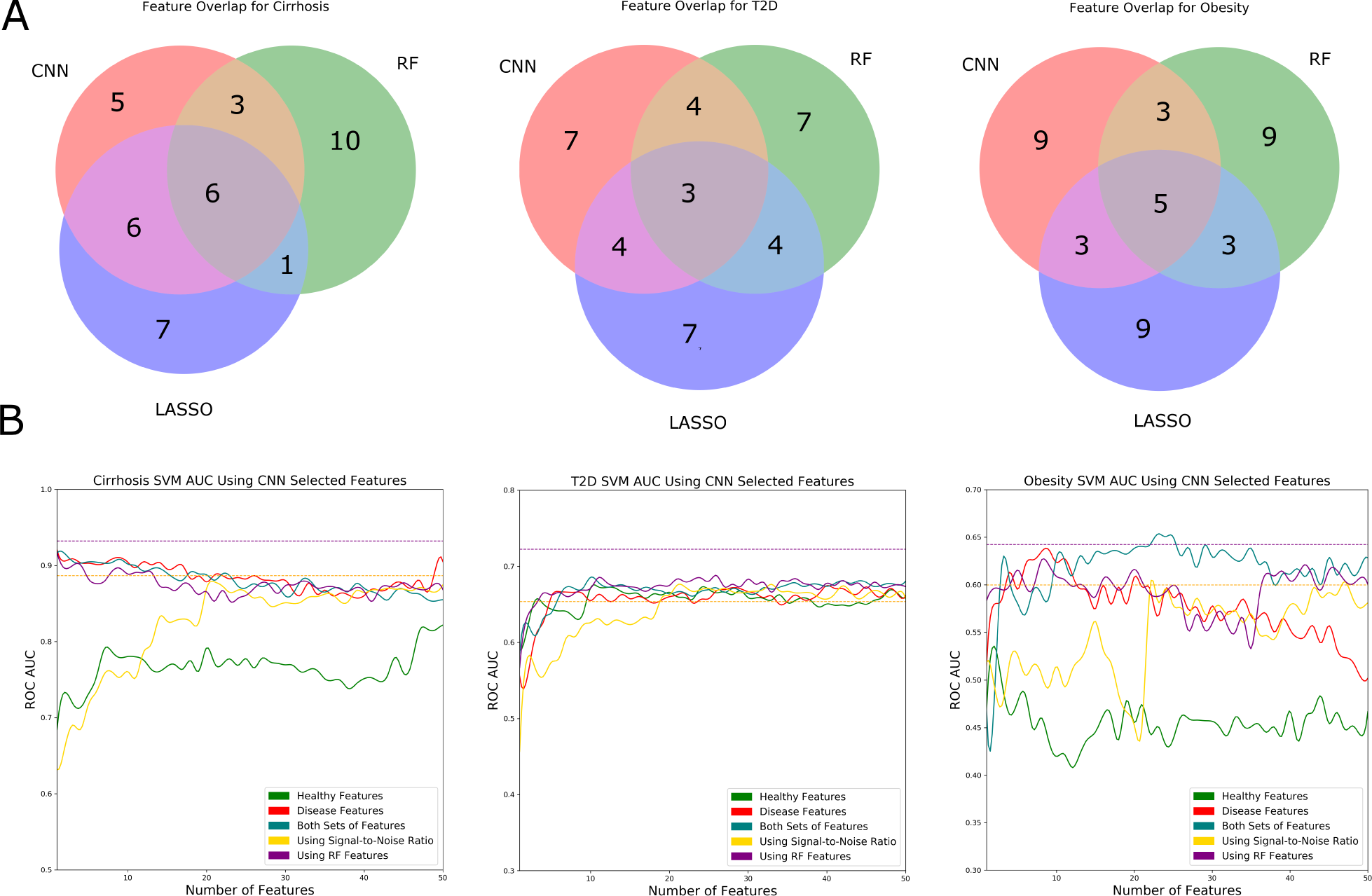
Benchmarking of feature selection using features extracted from PopPhy-CNN. (A) Venn diagrams of the top twenty features selected from the combined set of features from PopPhy-CNN, RF, and LASSO. (B) Features extracted from PopPhy-CNN are benchmarked against features found by RF (purple) and by using signal-to-noise ratio (yellow). Three different sets of features from the PopPhy-CNN are represented: disease (red), healthy (green), combined set (teal). The selected features are used to train an SVM model using 10 times 10-fold cross validation. The orange horizontal dashed line is the average AUC computed for SVM using the whole feature set and the dashed purple line is the average AUC for RF using the whole feature set.

Next we evaluated whether the sets of features could be used in building better prediction models in SVM, which is the only model that does not have any feature selection capacity in our evaluation. To do this, we trained SVM models using the top features ranging from 1 to 50. For PopPhy-CNN, we again only use OTUs found in the original OTU vector. We also combined the features from both classes into a single ranked list for benchmarking. For a baseline comparisons, we used a ranking list based on signal-to-noise ratio as well as the average feature rankings from the RF models. The SVM models were trained using 10 times 10 fold cross-validation using the same partitions and trained in the same way as described in the model evaluation. The results for all the three feature ranking sets are summarized in Fig. 7(B). Again, we observe that cirrhosis is best predicted by the OTUs found important in the cirrhosis patients, as the set of important OTUs in cirrhosis patients performs the best. For the T2D dataset, all models performed about the same, which may imply that there may have been only a few discriminatory features that were easy to capture, but these features could only predict a portion of the dataset well. In the obesity dataset, we observe a distinct pattern. Neither the healthy nor the disease features perform well on their own. However, when they were combined, we see a significant improvement in AUC. The resulting SVM models even outperformed the original RF model at a few points.

In a separate study in which we only selected the top 10% of maximum activations and the top 10% most frequent activation locations, we did not notice a difference in the performance of the cirrhosis features or the type 2 diabetes features. However, the obesity features were not able to improve SVM at all and both sets as well as the combination set of features oscillated around 0.5. When we removed the filtering, we noticed a change in the features selected in the obesity dataset as well as the improved performance.

Taken together, the evaluation demonstrates that our feature extraction procedure can identify more informative features than traditional feature selection methods. Using these features in SVM models led to improved performance with the best improvement observed in the hardest dataset.

### 3.6 Computation time

The PopPhy-CNN was implemented using the Theano library and was run using an NVIDIA Titan XP GPU. Table 2 shows the average time for one network to train. We trained each network 400 epochs using stochastic gradient descent (batch size of 1).

**Table 2.**
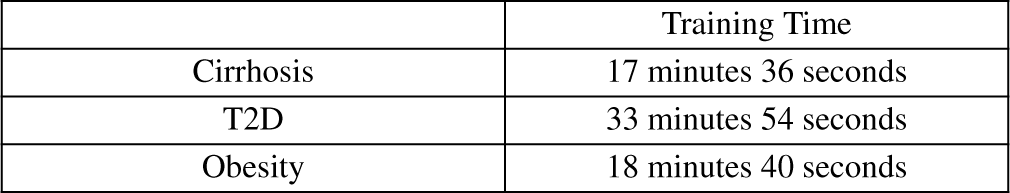
Training Time

## 4 Discussion

We have developed a novel architecture for CNN, PopPhy-CNN, for the prediction of the host phenotype from a metagenomic sample. The key contribution is leveraging biological knowledge in microbial taxa relative abundance profiles through a phylogenetic tree by our novel propagation and embedding procedure. The 2D matrix input obtained from this procedure enables CNNs to exploit the topological structure of the phylogenetic tree for developing more accurate predictive models. Using three metagenomic datasets, we showed that PopPhy-CNN models are more accurate than the model with the conventional vector input (1D-CNN), which does not take advantage of the biological knowledge in the phylogentic tree. In addition, we also shown that PopPhy-CNN outperformed RF, SVMs and LASSO models, establishing the evidence that CNNs can deliver more robust performance without requiring excessively large training sets.

We have also demonstrated the feasibility of extracting informative features from the learned CNN models and showed that the extracted features can improve the performance of SVMs compared with the models built on the entire feature sets. In addition, SVMs with the selected feature sets also performed better than SVMs trained on features ranked based on the criterion of the signal-to-noise ratio and features taken from RF models. This is especially intriguing, because the results provide the evidence that the activation maps on the first layer of the CNNs maintain spatial relationship between the microbial taxa on the phylogenetic tree. This implies that PopPhy-CNN benefits from learning informative features on the populated phylogentic tree embedded in the matrix format.

The use of phylogentic tree to imprint relevant biological knowledge in metagenomic data has been seen in several different machine learning models. For example, a class of phylogenetic-based feature weighting algorithms was proposed to group the relevant taxa into clades and the highly ranked groups in conjunction with RF had an improved classification performance [30]. The phylogentic information was also utilized in sparse linear discriminant models with the simultaneous use of intermediate nodes and leaves on a phylogentic tree [31].

There are a few applications using DNN for host phenotype prediction. The first large scale evaluation is the application of multilayer and recursive neural networks (RNNs) to determine body parts where the microbiome samples were taken using the input form of OTU vectors [11]. It has shown different performance of DNN models; a simple layer neural network and RF performed better than DNN models, and that RNNs could reveal a hierarchical structure among the samples. However, CNNs were not used in their study, and no explicit biological knowledge represented by the OTUs were explored in other DNNs for model learning. In a recent preprint, Fioravanti *et al*. described a different CNN architecture that explores the distance between nodes on a phynogentic tree by the patristic distance (the sum of the lengths of all branches connecting two OTUs on the tree) [29]. Their approach is to embed the phylogenetic tree in an Euclidean space. The computational analysis reported promising results on a synthetic data and on a gut metagenomics data from 222 inflammatory bowel disease patients and 38 healthy subjects, compared to linear SVMs, RF and a baseline fully connected multi-layer perceptron neural network.

The development of techniques to interpret CNNs models and using CNNs for extracting deep features have been an active research topic. For example, one study shows that a pre-trained CNN can be used for extracting local features based on one dataset, and retrained for classification for distinct datasets [32]. In another study, a novel CNN architecture is proposed to learn low dimensional CNNs for image retrieval in high-resolution remote sensing [33]. In addition, CNNs are applied for EEG decoding and for visualizing the informative EEG features [34]. In a recent paper, Shrikumar and colleagues proposed an algorithm that promotes the learning of important features through propagating activation differences [35]. It presents a successful example in identifying regulatory DNA motifs.

There are several directions for further study. The phylogenetic tree is the one of the core components in the PopPhy-CNN learning framework. It is likely that different trees constructed from different methods may affect the predictive performance and may also identify different microbial features. Furthermore, the current embedding scheme is designed to prevent sparsity in the matrix while preserving spatial phylogenetic relationships. This can create areas where the descendant nodes are not directly under their ancestors, allowing for unique patterns to be picked up by the CNN. However, one shortcoming arises when descendant nodes are shifted far enough away from the ancestral nodes, preventing CNN kernels from capturing them together. Therefore, different ways of embedding the populated trees into the matrix format may also affect the model performance.

Although the evaluation of our CNNs was conducted using the OTUs profiles obtained from the WMS sequencing platform, our method can be readily applied to the metagenomic data represented at any taxonomic levels if the number of taxa and the sample size are in the similar magnitude as the ones used in this work. However, if the number of microbial taxa substantially outnumbers that of the learning samples, more effective regularization schemes or algorithms that promote the learning of important features in CNNs are likely necessary.

## 5 Conclusion

PopPhy-CNN can be readily used for developing a predictive model from a metagenomic dataset of moderate size. It also facilitates the extraction and visualization of a ranked microbial taxonomic set for biological interpretation of the learned predictive model.

## Acknowledgements

We gratefully acknowledge the support of NVIDIA Corporation with the donation of the Titan Xp Pascal GPU used for this research.

